# Root Hydraulic Conductivity and Transpiration in Arabidopsis: Coordination Revealed by a High-Stomatal-Density Mutant

**DOI:** 10.64898/2025.12.17.694893

**Authors:** Pablo Daniel Cáceres, Carlos Augusto Manacorda, Moira Romina Sutka, Sebastián Asurmendi, Gabriela Amodeo, Irene Baroli

## Abstract

**Background and aims:** Understanding the coordination between root and shoot hydraulics is fundamental for improving plant performance under water stress. In this study, we investigated how shoot traits that enhance transpiration influence root hydraulic properties, using the *Arabidopsis thaliana* double mutant *epf1 epf2*, characterized by high stomatal density and increased transpiration.

**Methods:** Plant lines *epf1 epf2* and Col-0 (wild type) were grown hydroponically and compared for stomatal traits, rate of water loss, leaf and root water relations, aquaporin expression, and root hydraulic conductivity (Lp_r_). Then, to assess responses to water deficit, osmotic stress was induced by adding 2% polyethylene glycol (PEG) to the nutrient solution seven days before measurements.

**Key results:** The *epf1 epf2* double mutant exhibited ∼150% higher stomatal density, yet stomatal conductance and short-term rosette water loss increased by only ∼30% relative to wild type. Despite higher water loss, the mutant maintained its leaf relative water content, concomitant with a more negative leaf osmotic potential; root osmotic potential was similar between genotypes. *epf1 epf2* showed lower Lp_r_ than Col-0. Aquaporin transcript levels and the relative aquaporin contribution to root water transport did not differ between genotypes. Under osmotic stress, Col-0 instead showed lower Lp_r_ than *epf1 epf2*, again without changes in aquaporin expression or relative contribution.

**Conclusions:** Our results highlight an active contribution of the root as a modulator of the whole-plant hydraulic balance. Across scenarios where xylem tension was expected to increase, stomatal aperture and Lp_r_ decreased. We suggest that enhanced transpiration elevates xylem tension, which acts as a long-distance cue, eliciting coordinated reductions in stomatal aperture and Lp_r,_ thereby constraining water flux.

## Introduction

The increased frequency of droughts and floods caused by climate change is reshaping terrestrial ecosystems and posing significant challenges to plant productivity (Nolan et al., 2018). Addressing these constraints requires the development of sustainable strategies to enhance plant resilience, which in turn demands a deeper understanding of how plants manage their water status. While traditional research on plant hydraulics has emphasized stomatal control of transpiration (Sack and Holbrook, 2006; Buckley, 2019), more recent findings highlight the critical roles of water uptake capacity and root hydraulic conductivity (Lp_r_) as key regulatory nodes in plant water relations (Vadez, 2014; Maurel & Nacry, 2020). Therefore, elucidating the basis of the hydraulic coordination between shoot and root is critical for optimizing water use and guiding breeding programs for improved crop performance under drought (Bartletta et al., 2016).

Plant growth and development are strongly influenced by water status (Torres-Ruiz et al., 2023). Water transport from roots to leaves depends on soil moisture, plant hydraulic traits, and atmospheric demand (Passioura, 1982; Boyer, 1985). Plants respond to the external constraints of soil conditions and climate by modulating their hydraulic properties, in the long term through structural changes, like root system remodeling (Dinneny, 2019; Maurel and Nacry, 2020) and stomatal patterning (Hetherington and Woodward, 2003; Zoulias et al., 2018). On shorter timescales plants regulate stomatal aperture and tissue hydraulics to maintain water balance and limit xylem cavitation (Javot and Maurel, 2002; Almeida-Rodriguez et al., 2011).

Shoot–root hydraulic communication is essential for whole-plant water homeostasis. Environmental perturbations affecting either organ can induce systemic responses in the other, mediated by signals that integrate changes in water status across tissues. For example, water stress, whether due to drought or waterlogging, generates root-derived signals, such as an increased abscisic acid content, which regulate stomatal aperture in leaves (Davies et al., 2005; Christmann et al., 2007). Conversely, changes in vapor pressure deficit or leaf shading can influence Lp_r_ by modifying apoplast size or membrane permeability (Sakurai-Ishikawa et al., 2011; Vandeleur et al., 2014; Meng et al., 2016).

Water flow in plants can be described by an Ohm’s law analogy: J = L·ΔΨ_w_, where L is the equivalent hydraulic conductance of the pathway considered and ΔΨ_w_ is the water potential difference between soil and atmosphere (Suku et al., 2014). This equation can be formulated for any pair of points within the plant by redefining the ΔΨw between those points. In roots, water uptake and transport can be explained by the combination of a radial and an axial component. Radial conductance is the primary limiting factor for water uptake (Steudle & Peterson, 1998; Frensch & Steudle, 1989). It comprises an apoplastic pathway (through cell walls and intercellular spaces) and a cell-to-cell (transmembrane and through plasmodesmata) pathway occurring through membranes and plasmodesmata (Steudle, 2000). In contrast, axial conductance refers to xylem-mediated transport by bulk flow (Steudle and Peterson, 1998).

Aquaporins are membrane proteins which significantly impact the permeability of the plasma and internal membranes of cells (Chaumont and Tyerman, 2014). Arabidopsis contains 35 aquaporin genes and 13 of them belong to the plasma membrane intrinsic protein (PIP) subfamily, which modulate water transport (Johanson, 2001; Perez Di Giorgio et al., 2014). Aquaporin activity can be rapidly regulated through changes in gene expression, phosphorylation, vacuolar internalization, cytosolic pH shifts, or calcium signaling (Bellati et al., 2010; Moshelion et al., 2015; Vitali et al., 2015; Ozu et al., 2022). Short-term changes in cellular water transport are primarily driven by the regulation of aquaporin expression and activity (Suku et al., 2014). Their contribution to root water transport in different arabidopsis accessions ranges from 30% to 77% of the total flow (Sutka et al., 2011). Diverse experimental studies conducted in various plant species have suggested that changes in transpiration at the leaf level can modulate the cell-to-cell pathway in roots by altering root aquaporin expression (Levin et al., 2009; Almeida-Rodriguez et al., 2011; Laur and Hacke, 2013; Liu et al., 2014; Vandeleur et al., 2014). Thus, aquaporins have been proposed as effectors of root-shoot hydraulic communication (Maurel et al., 2010).

Using Ohm’s law analogy, transpirational water flux through stomata can be expressed as the product of conductance and the driving force provided by the water vapor gradient between the atmosphere and the internal air spaces of the leaf. Stomatal conductance (g_s_) depends on the anatomical properties of individual stomata (their size, aperture and pore depth) as well as their number, so-called density (Dow et al., 2014; Franks et al., 2015; Nguyen et al., 2023). This dependence of transpiration rate on stomatal density (D) makes plants with altered D valuable tools to study shoot–root hydraulic coordination. The genes that control stomatal development have been extensively studied in arabidopsis, and several lines with altered D have been described (Zoulias et al., 2018; Bertolino et al., 2019), including the *epf1 epf2* in which the genes EPIDERMAL PATTERNING FACTOR 1 (EPF1) and EPIDERMAL PATTERNING FACTOR 2 (EPF2) are inactivated. These genes encode secreted peptides that regulate stomatal spacing and guard cell differentiation, and their combined inactivation leads to a 2.5-fold increase in D, higher transpiration rates, reduced biomass, and smaller leaf area (Doheny-Adams et al., 2012). Whereas these types of plants have been extensively studied for the impact of D on gas exchange (Dow et al., 2014a; Dow and Bergmann, 2014; Franks et al., 2015). Studies on their root phenotype are scarce and show contrasting results. Two reports linked higher D and transpiration rate with larger root systems and enhanced nutrient uptake (Hepworth et al., 2015; Hepworth et al., 2016), while another found no differences in biomass or root architecture in the *epf2* single mutant (Mawodza et al., 2022). However, none of these studies have addressed the root functional properties of high D mutants, a task that requires hydroponic systems to enable precise physiological measurements.

In this study, we investigated the relationship between the hydraulic properties of roots in the *epf1 epf2* arabidopsis double mutant, which exhibits a significant increase in D and transpiration rate (Dow et al., 2014b). We also examined how the mutant adjusted its water balance under mild osmotic stress, exploring the role of key water-transporting aquaporins in this process. We hypothesized that the increased transpiration in *epf1 epf2* would drive compensatory adjustments in Lp_r_ to sustain water supply to the shoot under both control and osmotic stress conditions. These adjustments would rely on key regulations of the cell-to-cell water transport pathway through aquaporins. To our knowledge, this is the first study to explore how developmental differences impact the physiological coordination of water transport between root and shoot, shedding light on the dynamic regulation of hydraulic balance during vegetative growth. By contrasting the hydraulic conductance and water transport efficiency in mutant and wild-type lines, we aimed to explore the coordination between transpiration and root hydraulic properties during plant development.

## Materials and methods

### Plant material, growth conditions and osmotic stress treatment

Seeds of *Arabidopsis thaliana* Col-0 and the double mutant *epf1 epf2* (Doheny-Adams et al., 2012; kindly provided by Prof. Julie Gray, University of Sheffield, UK) were sterilized and sown on Murashige and Skoog (MS) medium (Murashige & Skoog, 1962) stratified at 4°C for 2 days and transferred to a growth chamber with a temperature of 21-22°C, a relative humidity of 55-65% and a short-day photoperiod (10/14). The photosynthetically active radiation (PAR) was 115 (±10) μmol m^−2^ s^−1^. To facilitate better access to the roots, plants were grown in a hydroponic system. A nutrient solution with approximately osmotic (water) potential of -0.11 MPa, adapted from Conn et al. (2013), was continuously aerated using an aquarium pump and replaced weekly. Measurements were taken 36 days after germination. To induce osmotic stress, a 2% (w/v) polyethylene glycol (PEG 6000) solution was prepared using the nutrient solution as the solvent, with a final osmotic potential of approximately -0.24 MPa. For osmotic stress treatment, 2% PEG was applied 7 days prior to measurement, while control plants remained under standard conditions.

### Stomatal density, initial rate of water loss, stomatal conductance, and maximum stomatal conductance

D was determined as the number of stomata per unit leaf area from imprints of the abaxial epidermis (Baroli et al., 2008). The same plants were used to measure stomatal conductance (gₛ) and root water management parameters. In all cases, gₛ was measured using a steady-state porometer (SC-1, Metergroup, Pullman, WA) on the abaxial side of the youngest fully expanded leaf, keeping plants under their growing conditions; measurements were taken between ZT1 and ZT7 (ZT0 = lights on). The maximum stomatal conductance (gₘₐₓ) was estimated based on anatomical parameters obtained from a sub-set, randomly chosen, of the same epidermal imprints used to determine D, according to the following formula:

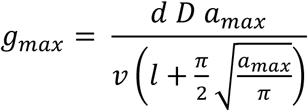

where a_max_ (maximum stomatal pore area) 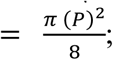 p = stomatal pore length; l = mean pore depth; d = diffusivity of water vapor in air and v = molar volume of air (Dow and Bergmann, 2014). The initial rate of water loss of the whole rosette was measured gravimetrically. Excised rosettes were immediately placed, abaxial side up, on an analytical scale, and weight loss was recorded every 15 seconds for 180 seconds, rate of water loss was then calculated as the slope of a linear equation fit of the initial points (R^2^>0.98).

### Osmotic potential and relative water content

The osmotic potential of the youngest fully expanded leaf and the first centimeter of the main root apex was measured using a vapor pressure osmometer (Vapro 5520, Wescor, USA) according to Sutka et al. (2016). The relative water content (RWC) of the youngest fully expanded leaf was determined as by Sutka et al. (2016).

### Root hydraulic conductivity and percentage of aquaporin contribution

Lp_r_ was measured in de-topped plants using a Scholander pressure chamber (Model 4, BioControl, Argentina). The root system was placed in a 50 mL Falcon tube with culture solution and inserted into the chamber. The hypocotyl was attached to a glass capillary (internal diameter, 0.86 mm), sealed with dental paste (A+ Silicone, Densell; Argentina). Roots were successively subjected to 0.20, 0.30 and 0.40 MPa and the exudate flow rate (J_v_) was obtained from meniscus displacement in the calibrated capillary, creating a flow curve to calculate hydraulic conductance (Lp = J_v_ / P). Data were normalized by root dry weight. Measurements were performed between ZT1 and ZT7 (ZT0 = lights on). Sodium azide, a commonly used pharmacological inhibitor of aquaporins (Sutka et al., 2011; Shahzad et al., 2024), was used to study the contribution of aquaporins to the observed Lp_r_. Briefly, plants were incubated in a solution with 5 mM KNO_3_, 2 mM MgSO_4_, 1 mM Ca(NO_3_)_2_, and 10 mM MES, pH 6, with 20 mM KCl for 20 minutes at 0.3 MPa, then treated with 1 mM NaN_3_. J_v_ (P) was measured at 0.3 MPa continuously until a steady low value was attained (typically about 20 min). Such value was taken to represent the maximum flow inhibition, and the remaining flow was considered to represent the aquaporin-independent contribution to the overall flow (Sutka et al., 2011).

### Root anatomical measurements and leaf and rosette area

To anatomically characterize the root and compute root and vascular cylinder diameter, freehand cross-sections were taken from the first centimeter of the root apex and analyzed using optical and epifluorescence microscopy (Zeiss Axioskop 2, Japan). Images were captured with a Nikon E8700 digital camera. The projected area of the rosette was measured by photographing freshly excised rosettes placed on a contrasting background and analyzing the images with ImageJ software (version 1.53t, http://imagej.nih.gov/ij).

### Gene expression analysis

The aquaporin genes for expression analysis were selected by comparison with publicly available arabidopsis gene expression databases (Genevestigator V3 (Hruz et al., 2008) and eFP-Browser, https://www.bar.utoronto.ca/). Focus was made on root and/or shoot organ specific expression and on expression that responded to water deficit stress. Oligonucleotide primer sets for quantitative real-time PCR were designed using Primer BLAST software (Ye et al., 2012) and are listed in Table S1. Details on the minimum information for publication of quantitative real-time PCR experiments (MIQE Guidelines; (Bustin et al., 2009)) are listed in Table S2. Primers were designed to target exon regions and all splice variants as predicted by ThaleMine (https://www.bar.utoronto.ca/). For qPCR analysis, LinReg PCR program (Ramakers et al., 2003) and the normalization method of (Pfaffl et al., 2002) were used. Relative expression ratios and statistical analysis were calculated using fg Statistics software (Di Renzo, 2009).

### Experimental design and statistical analysis

All measurements were performed at the same developmental stage and time window during the day (plants measured between ZT1–ZT7). For the PEG experiment, 2% PEG was applied 7 days before measurement; control plants from the same sowing cohort were measured under identical environmental conditions. All statistical analyses were performed using R (R Core Team, 2020; RStudio Team, 2019). Data normality and homoscedasticity were evaluated using Shapiro–Wilk and Levene’s tests, respectively. For each response variable we fitted linear (LM) and linear mixed-effects models (LMM; lme4::lmer) with Genotype (and Treatment where applicable) as fixed effects and experiment/batch as a random intercept. Competing models were compared by AIC and the best-supported model was used for inference. Statistical significance for fixed effects was evaluated with F-tests from an ANOVA on the final model. Results are reported as means with 95% confidence intervals; all individual data points are shown in the figures. Significance thresholds: NS p>0.05, * p≤0.05, ** p≤0.01, *** p≤0.001. Owing to the time-intensive nature of hydraulic measurements and to preserve sample size, we analyzed control and PEG cohorts with separate linear/mixed models and restricted statistical inference to within-condition genotype contrasts.

## Results

### A marked increase in stomatal density results in only moderate changes in water loss

We aimed to investigate the hydraulic coordination between roots and shoots in arabidopsis, by comparing the wild type (Col-0) and a genetic line within the same background, the *epf1 epf2* double mutant, which shows enhanced transpiration due to its increased D. The effect of the combined *epf1* and *epf2* mutations on D and size in plants grown in soil is well-documented (Doheny-Adams et al., 2012; Dow et al., 2014a). However, to quantitatively assess the hydraulic contribution of roots, we opted for hydroponic culture, which provides easy access to the root system and allows for the controlled addition of inhibitors and modulators of root water transport. In hydroponics, the *epf1 epf2* mutant exhibited a significantly higher D compared to Col-0, with an increase of more than 150% (p ≤ 0.001; Fig. 1a). In parallel, the double mutant shows no significant differences in stomatal dimensions, among them, pore length or guard cell width (Table 1). Consistently, the average maximum theoretical stomatal conductance (g_max_; Dow and Bergmann, 2014) was 655.9 and 1766.2 mmol H_2_O m⁻² s⁻¹ for Col-0 and *epf1 epf2*, respectively, representing a 169% increase (p ≤ 0.001, Fig. 1c). This closely mirrors the difference in D, which is expected, as D is the only anatomical parameter varying in the g_max_ equation. Despite the substantial increase in D, the *epf1 epf2* mutant showed only an approximately 30% increase in the measured stomatal conductance (g_s_) relative to Col-0, with values of 329.6 ± 14.8 mmol H_2_O m⁻² s⁻¹ for Col-0 and 421.3 ± 15.3 mmol H_2_O m⁻² s⁻¹ for *epf1 epf2* (p ≤ 0.001; Fig. 1d). Short-term (180 s) water loss of excised rosettes revealed a similar trend, with *epf1 epf2* plants losing water at a 22% faster rate than Col-0 under the tested conditions (p ≤ 0.05, Fig. 1b). These results indicate that while the *epf1 epf2* mutant displays a marked increase in D, its g_s_ short-term water loss are only moderately affected, even when growing under conditions of unlimited water supply.

**Figure 1.**
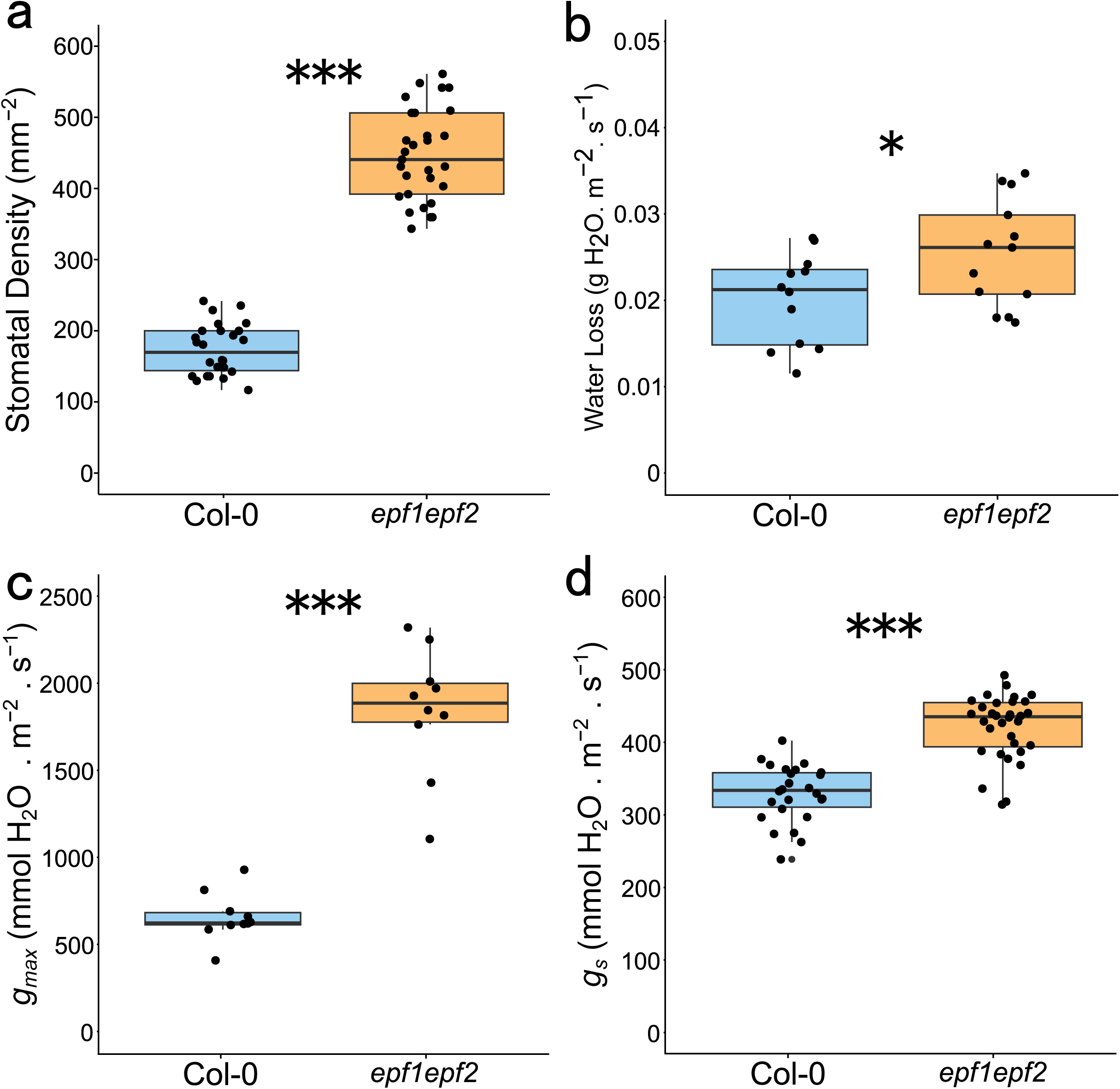
Impact of mutations in EPF1 and EPF2 on stomatal parameters and water use in hydroponically grown plants. (a) Stomatal density; (b) initial rate of water loss, measured gravimetrically during the first 180s after excising the rosette; (c) maximum stomatal conductance (g_max_); and (d) measured stomatal conductance (g_s_). Each dot represents an individual plant, and the boxplots show median (center line), interquartile range (box), and whiskers = 1.5×IQR. Significance: * = p ≤ 0.05, *** p ≤ 0.001. Combined results from three (a, c, d) or two (b) experiments, which yielded similar results, are presented.

**Table 1:**
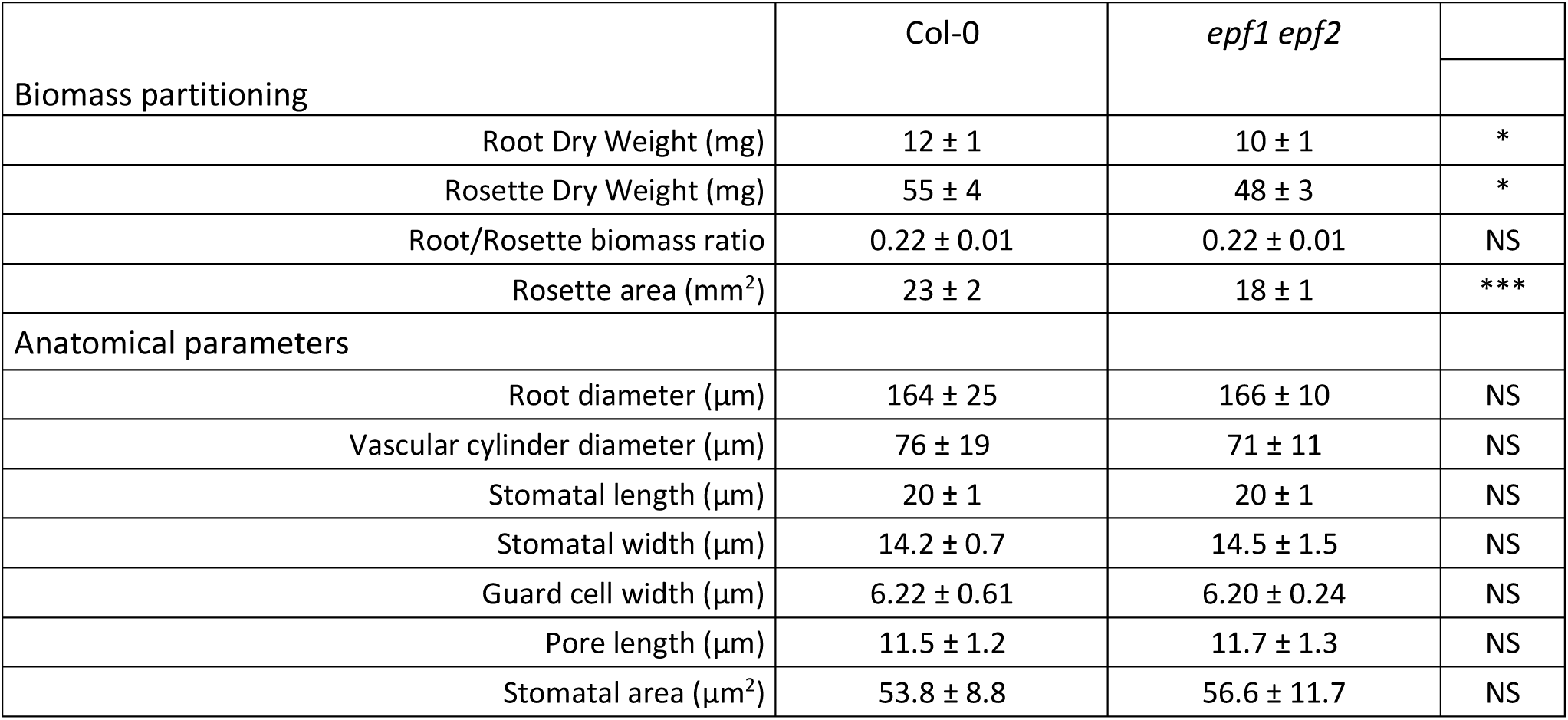
Plant biomass and anatomical parameters.

### Uncoupled osmotic responses between roots and leaves in *epf1 epf2* mutants

To analyze the water status at tissue level we quantified the leaf relative water content and the osmotic potential of leaves and roots. We observed a mild and non-significant decrease in leaf RWC in *epf1 epf2* plants (Fig. 2a), whereas their leaf osmotic potential was significantly lower than in Col-0 plants (p ≤ 0.001, Fig. 2b). In the root, the osmotic potential did not differ significantly between genotypes (Fig. 2c). These results indicate that there are no changes in root osmolytes coupled to the significant changes in leaf water status.

**Figure 2.**
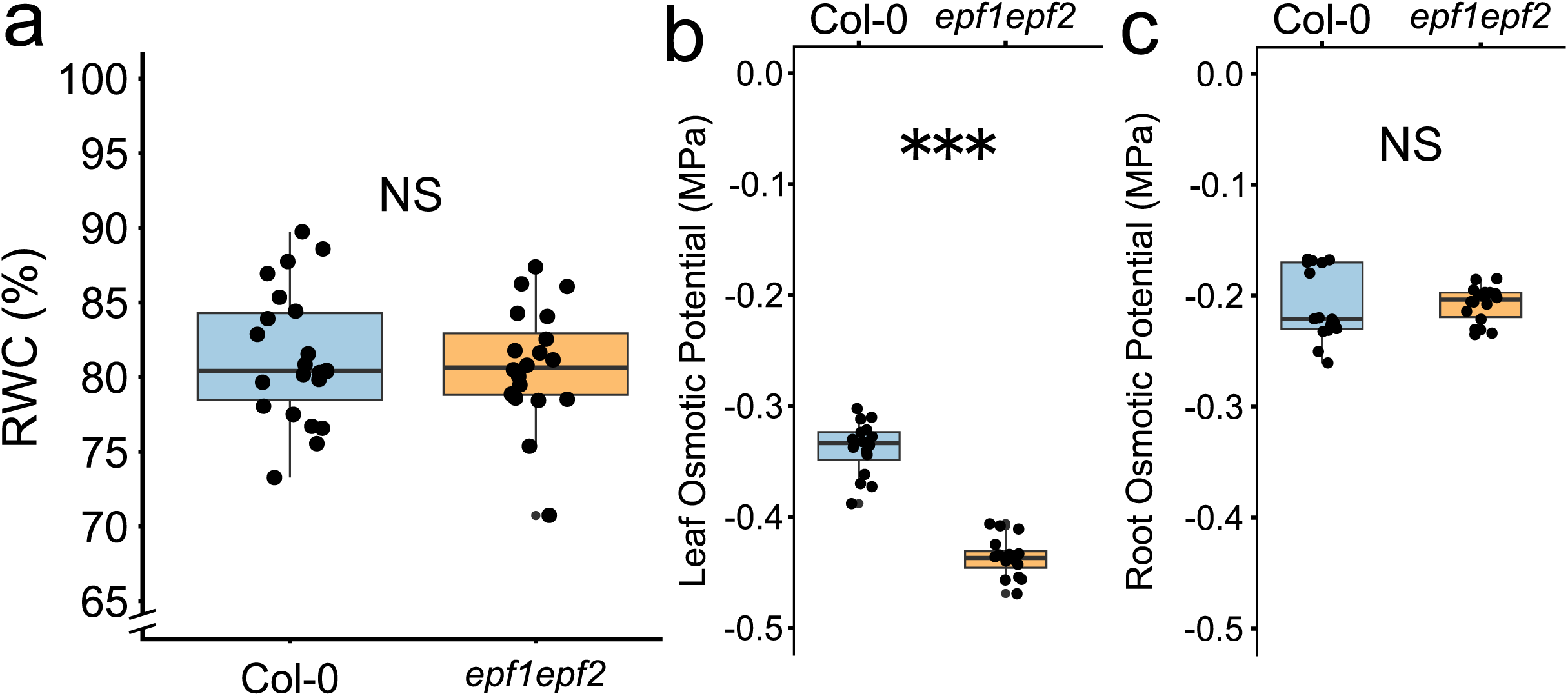
Water status parameters in *epf1 epf2* and wild-type plants. (a) Leaf relative water content (RWC), a broken y-axis was used to better visualize the differences; osmotic potential in (b) leaves, and (c) roots. Each dot represents an individual plant, and the boxplots display the median (center line) and interquartile range. NS = p > 0.05, ** = p ≤ 0.01; Data were obtained from two independent experiments which yielded similar results.

### Biomass and anatomical parameters in *epf1 epf2* mutants

Hydroponic growth provided a controlled framework to evaluate whether the increased transpiration of the *epf1 epf2* mutant leads to compensatory changes in root development or anatomy parameters that have not been previously characterized in this mutant. Our results are shown in Table 1. Both root and rosette dry weights showed a slight but significant decrease in *epf1 epf2* plants compared to Col-0, while the root/rosette biomass ratio remained unchanged. The rosette biomass decrease is consistent with previous findings in soil-grown plants (Doheny-Adams et al., 2012), suggesting that this reduction is a robust feature of the mutant across growth conditions. Among the analyzed traits, rosette area was the most affected by the mutations, without observable differences in leaf morphology and trichome distribution. To explore whether altered transpiration affects radial water transport capacity through structural changes, we analyzed root anatomical traits associated with hydraulic resistance. Both genotypes had similar root and vascular cylinder diameter (Table 1). No observable differences were detected between genotypes under fluorescence microscopy of transverse histological sections stained with Sudan IV, which allows for the visualization of suberin deposition in the endodermal Casparian strips and provides information on the barrier regulating water movement through the apoplastic pathway (data not shown).

### Plants with moderately increased transpiration rate have a lower root hydraulic conductivity without impacting relative pathway distribution

We evaluated the impact of an increased transpiration on Lp_r_ under the unrestricted water supply afforded by hydroponic culture. Despite their increased transpiration, *epf1 epf2* plants exhibited nearly a 15% decrease in Lp_r_ compared to Col-0 (p ≤ 0.01; Fig. 3a). We assessed the aquaporin contribution of the cell-to-cell pathway using sodium azide, as a inhibitor of aquaporins (Sutka et al., 2011). In both genotypes sodium azide inhibited nearly 80% of the total water flow, indicating a similar relative aquaporin contribution (Fig. 3b). However, the absence of significant differences between genotypes suggests that the reduction in Lp_r_ observed in *epf1 epf2* may result from a proportional decrease in both the cellular and apoplastic pathways, thus maintaining the relative contribution of each route to the total flow. Expression analysis of the 13 major water-transporting aquaporins in roots and rosettes revealed no significant differences in transcript levels between genotypes (Fig. 3c).

**Figure 3.**
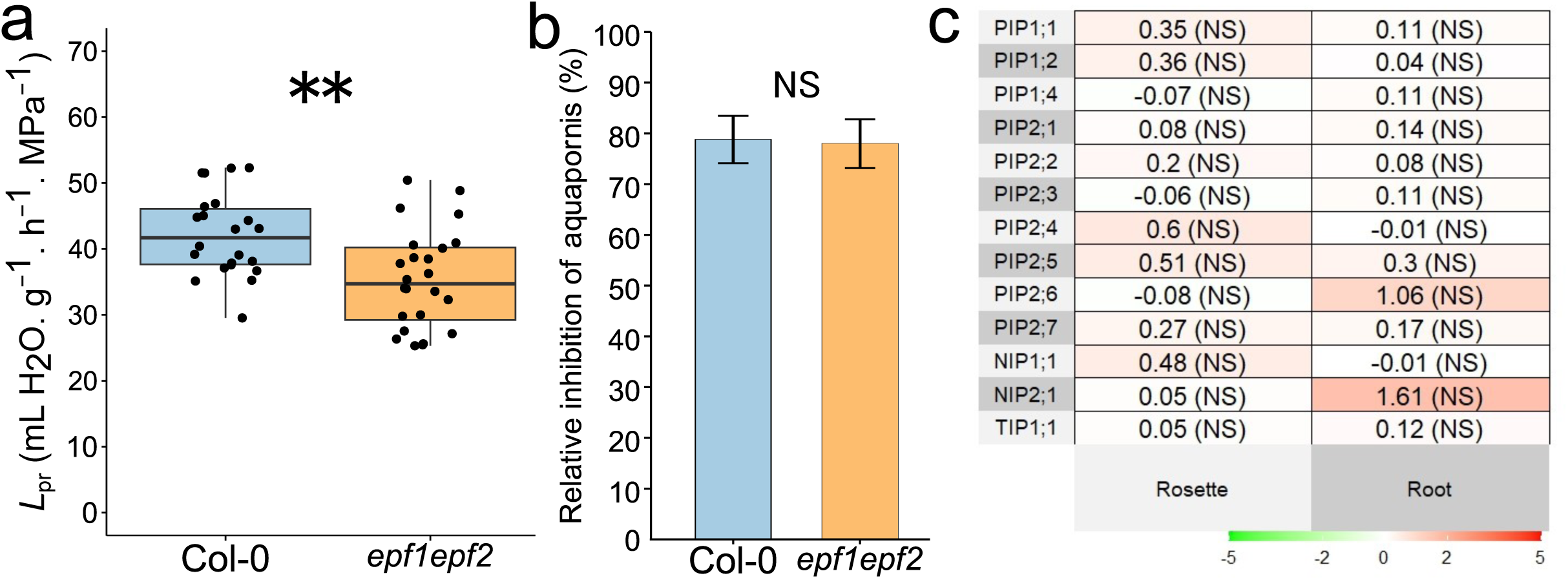
Root water management in wild type and *epf1 epf2* plants. (a) root hydraulic conductance normalized by root dry weight (Lp_r_); (b) relative inhibition of aquaporins by sodium azide treatment. Combined results from three experiments, which yielded similar results are presented. In (a) each dot represents an individual plant, and the boxplots show median (center line), interquartile range (box), and whiskers = 1.5×IQR. ** = p ≤ 0.01. (c) Relative expression analysis of aquaporin genes measured by qRT-PCR. The heatmap shows results for rosettes and roots. Col-0 plants were used to relativize aquaporin expression levels for *epf1 epf2* mutant plants. Log2 fold-change values are given in a double-color scale (green = repression; red = induction). Genes were differentially expressed between genotypes if they simultaneously had a difference in expression of at least 2 times in a linear scale and a p-value lower than P ≤ 0.05 after false discovery rate correction (Benjamini-Hochberg). No differentially expressed genes between genotypes were found following this criterion.

### Sustained root hydraulic conductivity despite prolonged osmotic stress in *epf1 epf2*

To induce a controlled stress that would prompt hydraulic adjustments in arabidopsis, we added 2% PEG to our hydroponic solution. This approach allowed us to evaluate whether the *epf1 epf2* mutant can modulate its water transport mechanisms under conditions of reduced water availability. The PEG concentration was selected based on preliminary trials (data not shown), as it generated a visible but moderate stress, while still allowing the plants to survive beyond one week, a period in which developmental or acclimatory adjustments are expected to emerge (Kreszies et al., 2018; Robin et al., 2021), rather than focusing solely on acute responses. After exposure to PEG, g_s_ of the youngest fully expanded leaf decreased in both genotypes in the short- and long-term (Fig. 4a), following similar overall kinetics. A significant reduction in g_s_ was observed at 4 hours for both Col-0 and *epf1 epf2* when compared to their respective controls, indicating a rapid stomatal response to osmotic stress. At 24 hours, g_s_ partially recovered in both genotypes, although values remained slightly lower in PEG-treated plants compared to controls, without reaching statistically significant differences. By day 7, Col-0 showed a significant decrease in g_s_ under osmotic stress, while in *epf1 epf2* it was also reduced but not significantly. Throughout the course, the difference in g_s_ between genotypes was consistently maintained, with the mutant exhibiting approximately 30% higher rate of water loss through the stomata.

**Figure 4.**
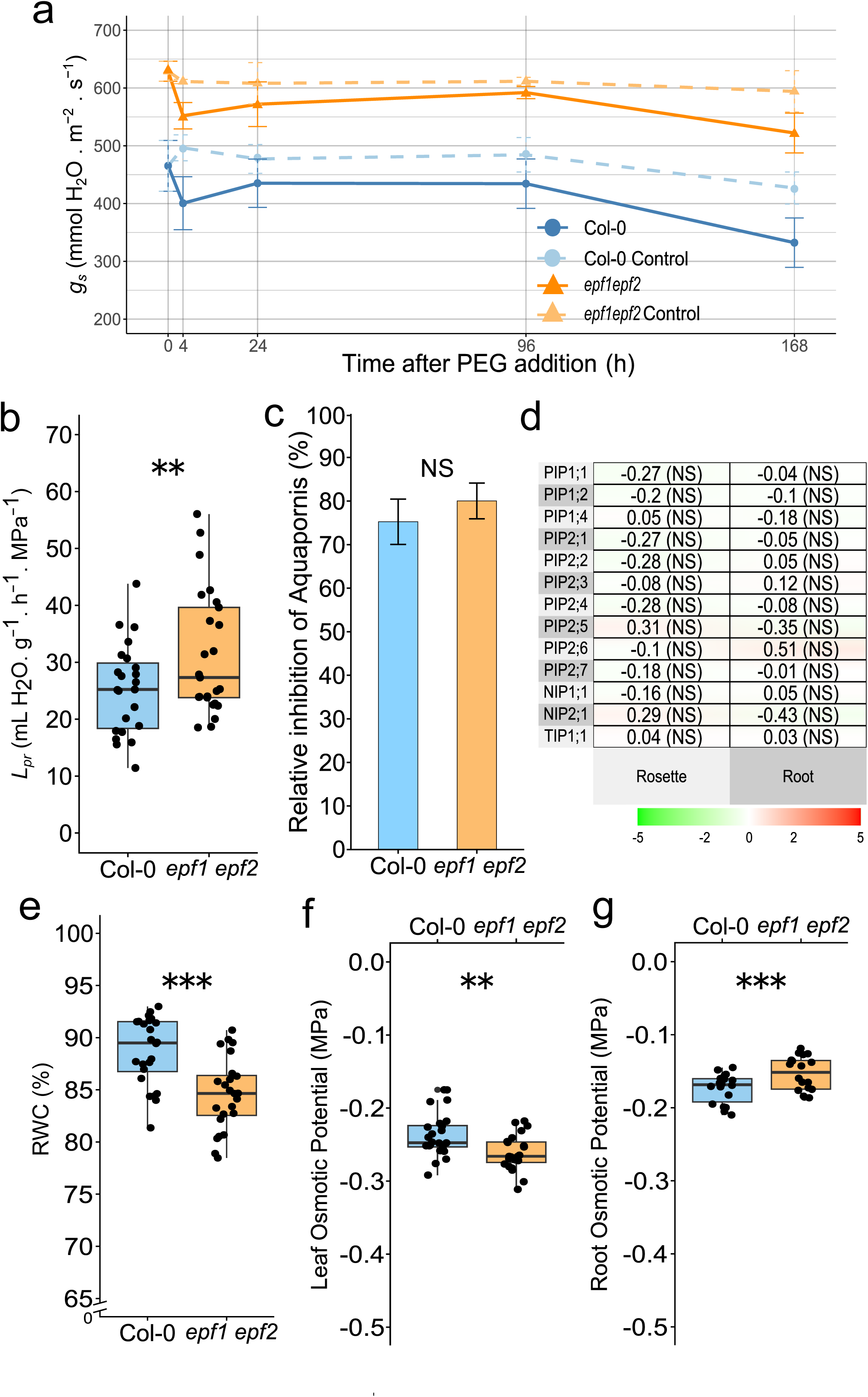
Hydraulic trait characterization of *epf1 epf2* and wild-type plants under osmotic stress. (a) Stomatal conductance (g_s_) during treatment with 2% PEG. Each data point represents mean values and error bars represent the confidence interval (p > 0.05). (b) Root hydraulic conductance normalized by root dry weight (Lp_r_); (c) relative inhibition of aquaporins by sodium azide treatment; (d) Relative expression analysis of aquaporin genes measured by qRT-PCR. Log2 fold-change values are given in a double-color scale (green = repression; red = induction). Genes were differentially expressed between genotypes if they simultaneously had a difference in expression of at least 2 times in a linear scale and a p-value lower than P ≤ 0.05 after false discovery rate correction (Benjamini-Hochberg). (e) leaf relative water content (RWC) a broken y-axis was used to better visualize the differences; (f) osmotic potential of leaves, and (g) osmotic potential of roots. (b-g) Hydraulic parameters measured after 168 h (7 d) incubation with 2% PEG. Each dot represents an individual plant, and boxplots show median (center line), interquartile range (box), and whiskers = 1.5×IQR. NS = p > 0.05, ** = p ≤ 0.01, *** = p ≤ 0.001. Combined results from two experiments, which yielded similar results are presented.

Under 2% PEG, Lp_r_ differed between genotypes, with *epf1 epf2* exhibiting higher values than Col-0 (33.2 vs. 25.9 ml H_2_O g^-1^ h^-1^ MPa^-1^; p ≤ 0.01; Fig. 4b). At the same time, a slight trend toward a higher relative contribution of aquaporin-mediated water transport was observed in *epf1 epf2* (80%) compared to Col-0 (75%), however, within-treatment differences were not statistically significant (Fig. 4c). Furthermore, qPCR analysis revealed no significant differences in the expression of the 13 major aquaporin isoforms known to be involved in root and shoot water transport (Fig. 4d).

To further characterize the impact of osmotic stress on plant water status, we also measured the RWC and osmotic potential of leaves and roots. RWC was significantly lower in *epf1 epf2* compared to Col-0 (84.6% vs. 88.7%, p ≤ 0.001; Fig. 4e), indicating a mild loss of tissue hydration in the mutant line. Leaf osmotic potential also showed a significant decrease in *epf1 epf2* (-0.26 MPa) compared to Col-0 (-0.24 MPa), suggesting a stronger osmotic adjustment in the mutant (p ≤ 0.01; Fig. 4f). Interestingly, root osmotic potential was slightly less negative in *epf1 epf2* (-0.15 MPa) than in Col-0 (-0.17 MPa; p ≤ 0.001; Fig. 4g), which may reflect differences in solute accumulation, water uptake capacity, or both, between genotypes under stress conditions.

## Discussion

In this study, we investigated shoot–root hydraulic coordination in arabidopsis by comparing wild-type and *epf1 epf2* plants, which differ in transpiration rates due to D but share similar morphology. We hypothesized that the fast-transpiring mutant would display compensatory adjustments in root hydraulic conductivity, requiring an active cell-to-cell pathway, to sustain water delivery under both optimal and osmotic stress conditions. This work presents the first comprehensive characterization of *epf1 epf2* hydraulics in hydroponics, assessing a wide range of physiological and anatomical traits. Comparison with previous soil-based studies of this mutant (Table 2) revealed consistent trends in D, stomatal conductance, and shoot biomass, validating hydroponics as a robust system to study *epf1 epf2*, which offers a unique advantage for investigating fully developed root systems in a controlled and homogeneous environment.

**Table 2:** Comparison of published data and the present results. . Approximate percentage change in key physiological and morphological parameters of the *epf1 epf2* double mutant relative to the wild-type line Col-0, across different studies and growth conditions.

| | Growing conditions | Stomatal density | Stomatal dimensions | | $g_s$ | $g_{max}$ | Rosette | | Root | |
| --- | --- | --- | --- | --- | --- | --- | --- | --- | --- | --- |
|  |  |  | Length | Area |  |  | Dry weight | Area | Dry weight | Area |
| Doheny-Adams et al., 2012 | Soil | +170% | n.d. | -50% | +60% | +170% | -10% | -20% | n.d. | n.d. |
| Dow et al., 2013 | Soil | +140% | -20% | -10% | n.d. | +80% | n.d. | n.d. | n.d. | n.d. |
| Franks et al., 2015 | Soil | +180% | n.d. | -30% | +60% | +130% | n.d. | n.d. | n.d. | n.d. |
| Hepworth et al., 2015 | Soil | +150% | n.d. | n.d. | +40% | n.d. | n.d. | n.d. | n.d. | n.d. |
| Hepworth et al., 2016 | MS-Sand | +90% | n.d. | n.d. | n.d. | n.d. | n.d. | n.d. | n.d. | +10% |
| This study | Hydroponics | +160%* | 0 (NS) | 0 (NS) | +30%* | +170%* | -10%* | -20%* | -20%* | n.d. |

In hydroponics, where water supply is ample, the large increase in D in *epf1 epf2* did not yield a proportional rise in g_s_ or initial rosette water loss, which remained only moderately elevated. This pattern is consistent with increased transpiration rate imposing a hydraulic constraint at the whole-plant level. We observed that *epf1 epf2* exhibited regulatory responses at both the stomatal and root levels as compensatory mechanisms to counterbalance the elevated evaporative flux. At the shoot, regulation occurred by limiting the stomatal aperture: g_s_ increased far below the 170% rise expected from gₘₐₓ (Fig. 1 and Table 2). When measured under the identical ambient conditions of growth, in which both genotypes are expected to have similar boundary layer resistance, Col-0, g_s_ was ∼50% of gₘₐₓ, but in *epf1 epf2* this percentage was only 23%, reflecting a tighter restriction. Thus, under ample water supply to the roots, increasing D does not automatically lead to proportionally higher g_s_. Operational constraints related to stomatal size, spacing, and guard cell physiology may limit this relationship, particularly when aperture is actively regulated in response to environmental or endogenous signals (Franks and Farquhar, 2007). Recent analyses confirm that operational leaf conductance depends equally on stomatal anatomy (D and size) and behavior (aperture), underscoring the joint structural and dynamic control of g_s_ (Ochoa et al., 2024).

Interestingly, despite their higher anatomical gas-exchange capacity, *epf1 epf2* plants were consistently smaller than Col-0, implying a cost of the altered stomatal patterning. Previous work has shown that g_s_ can increase markedly in *epf1 epf2* when exposed to low CO_2_ partial pressure in the gas-exchange cuvette, indicating that stomatal aperture in the mutant can approach its anatomical maximum when carbon availability becomes limiting. However, the responses observed in short-term gas exchange experiments may reflect the guard-cell autonomous, pore-specific response to intercellular CO_2_ levels (Dow et al., 2014a) and not the sustained behavior of stomata under steady state environmental conditions. In our system, with normal ambient CO₂ concentration and moderate humidity, stomata appeared finely tuned to restrict water loss despite high D, potentially at the expense of carbon gain and growth. Although D increased by ∼150%, g_s_ was only ∼30% higher compared to Col-0. The mutant did not fully compensate for the increased number of stomata by further stomatal closure, possibly because doing so would compromise CO₂ uptake and photosynthesis. These findings highlight that the D–g_s_–growth relationship is context-dependent, shaped by both anatomy and long-term regulation of water balance. Leaf RWC was similar between genotypes, indicating that *epf1 epf2* maintained leaf hydration despite increased water loss. However, *epf1 epf2* leaves showed more negative osmotic potential (Fig. 2), consistent with active osmotic adjustment to buffer turgor under stress (Blum, 2017).

Does the root provide the regulatory mechanism that constrains transpirational water flow in the double mutant? Previous studies reported a positive correlation between transpiration and root hydraulics when transpiration was experimentally reduced by drought or circadian rhythms. These responses were largely mediated by aquaporins and interpreted as evidence of a coordinated regulation between shoot water loss and root water uptake (Vandeleur et al., 2009; 2014; Sakurai-Ishikawa et al., 2011). In contrast, under hydroponics and with a high-transpiring mutant we found a reduced Lp_r_ (Fig. 3), in line with result of Meng et al. (2016), reinforcing the view that roots act as active rheostats modulating hydraulics with evaporative demand (Maurel et al. 2010) and thereby buffering whole-plant water balance in the face of anatomical alterations. In our experimental set-up, elevated transpiration was not linked to differences in expression of major aquaporins in either roots or shoots (Fig. 3). Sodium azide inhibition was consistent with a predominant transmembrane pathway. Interestingly, the relative contribution of aquaporins was similar between genotypes. Thus, the lower Lp_r_ in the mutant likely reflects a proportional decrease in both cell-to-cell and apoplastic pathways, preserving their relative contributions while reducing absolute flow. Given that aquaporin transcript levels were not altered, this evidence instead points to post-transcriptional mechanisms of regulation, consistent with the notion that aquaporins provide a sustained capacity for fine-tuning the cell-to-cell pathway (Maurel et al., 2015). Likewise, root osmotic potential did not differ, implying no osmotic adjustment at the root level and no evidence of hydraulic redistribution. As to the nature of the signal mediating the observed adjustments, while long-distance hormonal signaling (e.g. ABA routing) cannot be excluded, the non-limiting water availability in hydroponics would not favor a root-derived hydraulic stress signal; instead, the Lp_r_ reduction is best viewed as a response to a hydraulic cue (e.g. xylem tension) originated in the shoot.

To test whether stronger adjustments occurred under limited water supply, we imposed a moderate osmotic stress with 2% PEG. This method is widely used to simulate drought conditions in controlled environments without causing immediate damage (Verslues et al., 2006; Rosales et al., 2019). We selected a 7-day incubation period to allow for the establishment of long-term physiological adjustments to osmotic stress, such as endodermal suberization (Barberon et al., 2016), altered root architecture (Uga et al., 2013), or regulation of aquaporin expression (Ding et al., 2016). Both genotypes initially reduced g_s_ upon PEG treatment, but the extent of the difference in g_s_ between *epf1 epf2* and Col-0 observed under normal conditions persisted. After the early response, the mutant showed limited capacity to further reduce g_s_, remaining near control values (Fig. 4). This supports previous suggestions that high-D mutants, in which small guard-cell turgor changes produce larger variations in conductance, operate close to their g_max_, with reduced regulatory plasticity and lower water-use efficiency under stress (Doheny-Adams et al., 2012; Dow & Bergmann, 2014; Franks et al., 2015).

Under osmotic stress, the two genotypes exhibited contrasting root hydraulic responses. Whereas Col-0 showed a marked decline in Lp_r_, *epf1 epf2* showed no change, reversing the trend observed under control conditions (Fig. 3a and Fig. 4b). Although anatomical traits were not directly examined under PEG treatment, long-term osmotic stress is known to induce endodermal suberization and structural adjustments in arabidopsis roots (Barberon et al., 2016; Kreszies et al., 2018). However, the similar percentage of azide inhibition in both genotypes suggests that, if such anatomical modifications occurred, they were of comparable magnitude and did not differentially affect the relative contribution of the transport pathways. Expression of major aquaporins was similar between genotypes (Fig. 4). As under control conditions, the data point to post-transcriptional aquaporin regulation as a rapid means of controlling root water transport (Maurel et al., 2015). In *epf1 epf2*, the elevated transpirational demand associated with increased D may activate developmental feedback mechanisms that maintain transport capacity. Within the PEG treatment, leaf RWC was lower in *epf1 epf2* than in Col-0, indicating a mild reduction in tissue hydration in the mutant. Leaf osmotic potential followed a similar pattern, with *epf1 epf2* showing slightly more negative values. (Fig. 4). Taken together, the limited capacity of *epf1 epf2* to further close stomata or intensify osmotic adjustment explains the RWC decline, likely reflecting a ceiling in its hydraulic adjustment capacity. By contrast, wild-type plants minimized water loss through more effective stomatal closure and a reduction in Lp_r_. At the root level, osmotic potential was slightly less negative in the mutant, suggesting altered solute balance or cell volume regulation. This may reflect water partitioning between root parenchyma and the xylem that favors export to the shoot over local retention. Consistent with the stable Lp_r_ in *epf1 epf2*, this result supports the maintenance of radial water transport.

## Conclusions

Figure 5 summarizes and compares our data, in relative terms, with the expected outcome based on previous literature. We used g_s_ ^unit^= g /D as an estimation of unitary (per-stoma)_s_ stomatal aperture. We hypothesize that given the higher D of *epf1 epf2* and assuming that their g_s_ ^unit^ equals that of Col-0 under favorable water conditions, transpiration (E) should scale with D. Therefore, since the amount of transpired water must be incorporated by the roots (E ≈ J_v_), conceptually, two alternative scenarios could explain this coordination: (i) Lp_r_ increases to maintain a controlled xylem tension, or (ii) Lp_r_ remains unchanged, and xylem tension rises. Interestingly, neither scenario fully matched our observations, suggesting an alternative regulatory mechanism. The mutant behaved as if experiencing water stress, even under ample water supply, restricting g_s_ ^unit^ and Lp . This leads us to propose that increased xylem tension is the plausible signal triggering hydraulic adjustment. Under osmotic stress, wild-type and mutant plants exhibited contrasting strategies: *epf1 epf2* maintained high Lp_r_ despite reduced leaf hydration, whereas Col-0 showed lower conductance and tighter stomatal control. Overall, our results show that root hydraulics emerges as an actively regulated component of whole-plant water balance, modulated by shoot-derived cues rather than passively following transpiration demand. A plant with high evaporative capacity limits water flux by decreasing stomatal aperture and root hydraulic conductivity, likely as part of a coordinated adjustment to constrain water loss even under ample water availability. Using *epf1 epf2* as an experimental system reveals how persistent transpirational demand can be counterbalanced by the root: the cell-to-cell pathway acts as a rheostat, buffering whole-plant water flux, with the apoplastic route adjusting proportionally. Further investigation into the underlying mechanisms will be essential to advancing our understanding of plant water use efficiency and drought resilience.

**Figure 5:**
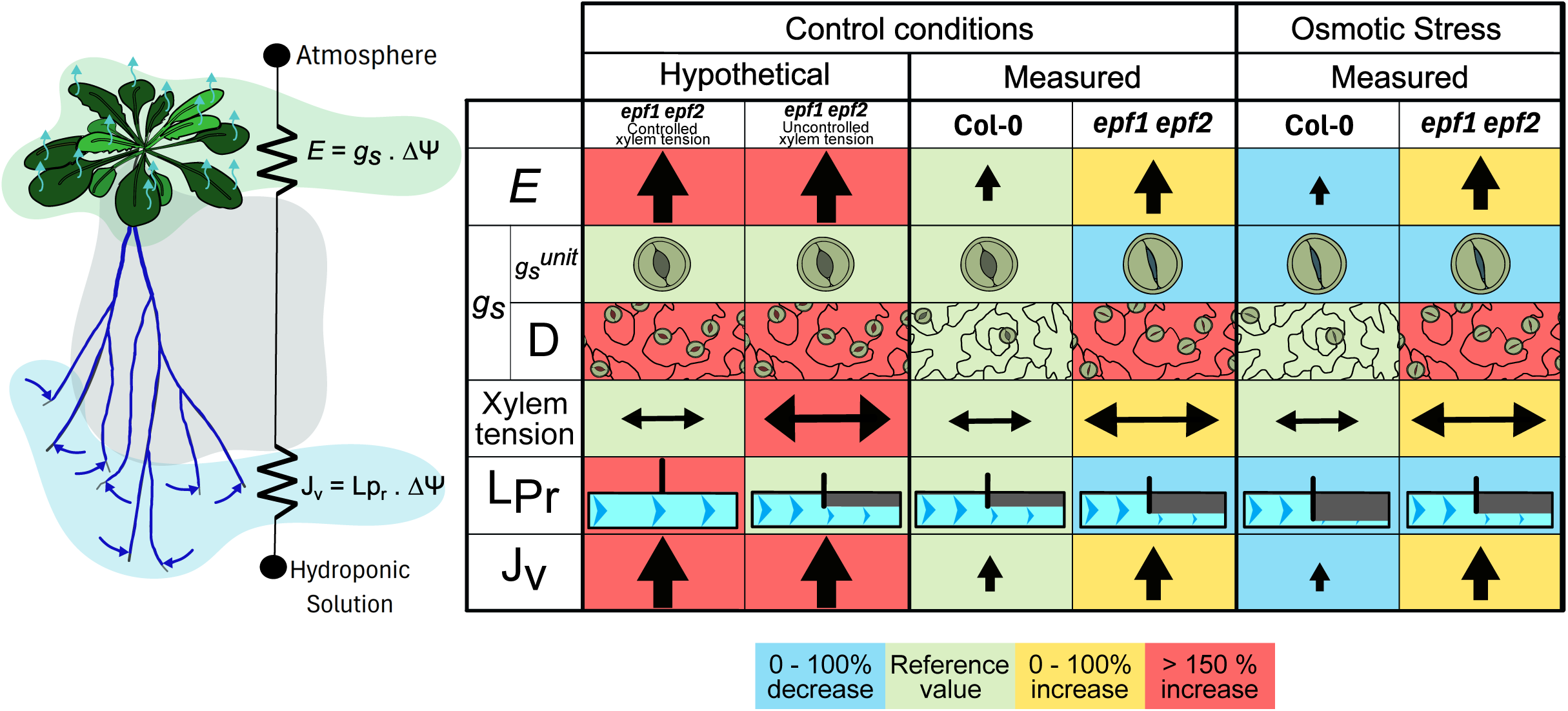
Summary model and conceptual synthesis of whole-plant hydraulics. The left panel shows a schematic model representation of arabidopsis hydraulic physiology, simplified as an analogy to an electric circuit with two conductances in series between two external points defining water potential gradient (ΔΨ). The table to the right summarizes expected (“Hypothetical”) and observed (“Measured”) changes under control and osmotic stress (2% PEG) in Col-0 and *epf1 epf2*. Two hypothetical alternatives are shown: controlled xylem tension (via higher Lp_r_) or uncontrolled xylem tension (with unchanged Lp_r_). Rows denote transpiration (E); stomatal conductance (g_s_); unitary stomatal conductance (g_s_ ^unit^= g /D as an estimation of per-stoma stomatal aperture.); stomatal density (D); xylem tension; root hydraulic conductivity (Lp_r_); and root water flux (J_v_). Single-arrows represent water movement; double-arrows represent xylem tension; in both, arrow thickness correlates with magnitude.

## Supporting information

Table Suppl 1 Oligonucleotide primers

Table Suppl 2 MIQE

## Acknowledgements

We thank Marina Recchi for expert technical assistance and Milagros Sampaolesi and Tomás Fiorini for help with plant culture. AI was used for help with English language editing. This work was supported by grants from the Agencia Nacional de Promoción Científica y Tecnológica, Argentina [PICT 2020-01438] to G.A. and [PICT 2021-697] to A. S.; from Instituto Nacional de Tecnología Agropecuaria [2023-PD-L01-I084] to A. S.; from Consejo Nacional de Investigaciones Científicas y Técnicas, Argentina [PIP 11220200101935CO] to I.B. and from University of Buenos Aires, Argentina [UBACyT 20020220100162BA] to I.B. None of the authors have a conflict of interest to disclose.

## Author contributions

P.D.C., C.A.M., S.A., G.A. and I.B. conceived and designed research. P.D.C., C.A.M., and M.S. performed the experiments. P.D.C. and I.B. prepared the manuscript. G.A., I.B. and S.A. obtained funding; all authors discussed the results and read and approved the manuscript.

