## Supplementary material for "Root Hydraulic Conductivity and Transpiration in Arabidopsis: Coordination Revealed by a High-Stomatal-Density Mutant": Table Suppl 2 MIQE

**Table S2.** Experimental conditions used for messenger RNA quantification by qRT-PCR assays based on MIQE requirements (Bustin *et al.*, 2009; Bustin *et al.*, 2010).

| **Experimental design** |  |
| --- | --- |
| Control groups | *Arabidopsis thaliana* Col-0 plants |
| Treatment groups | *Arabidopsis thaliana epf1 epf2* double mutant plants |
| **Sample** |  |
| Type of sample | Whole organ (rosette or roots) tissue for hydroponically grown short day conditions plants |
| Processing procedure | Liquid nitrogen homogenization |
| Sample frozen conditions | -80 ºC |
| Biological replicates | Four groups (Col-0 or *epf1 epf2* double mutants, rossette or root tissue). 3≤n≤6 biological replicates for each group |
| RNA: DNA-free | RT^-^ control without amplification |
| **RNA extraction** |  |
| Procedure | Acid Phenol extraction |
| Reagents | Transzol (TransGen Biotech) |
| Details of Dnase treatment | DNAse I Amp Grade (Invitrogen), 15 min at room temperature |
| Contamination assessment | < 0.09% (by means of RT- assessment) |
| Nucleic acid quantification | Absorbance at 260 nm |
| Instrument and method | NanoDrop instrument |
| Purity (A260/ A 280) | > 1.8 |
| Purity (A260/ A 230) | > 1.8 |
| RNA integrity | Analyzed by agarose gel electrophoresis |
| **Reverse transcription** |  |
| Complete reaction conditions | Reaction was performed as described by the manufacturer´s instructions |
| Amount of RNA and reaction volume | 1 µg of RNA, 20 µl |
| Priming oligonucleotide | Random primers (Invitrogen) |
| Reverse transcriptase | M-MLV (Invitrogen) |
| Temp and time | 10 min 50 ºC, 50 min 37 ºC, 15 min 70 ºC. |
| **qPCR protocol** |  |
| qPCR chemistry | SYBR green, ROX as passive reference. |
| Complete reaction conditions (*) | 5 min 94 ºC, (15 s 94 ºC, 30 s 60 ºC, 40 s 72 ºC) x 45 cycles |
| Reaction volume and amount of cDNA | 2 µl of a 1/20 dilution of synthesized cDNA in a final volume of reaction of 10 µl |
| Primers, Mg^2+^ and dNTPs concentration | 3 mM Mg^2+^, 200 nM primers, 0,2 mM dNTPs |
| Polymerase | Platinum® Taq DNA Polymerase (Invitrogen) |
| Buffer | 20 mM Tris-HCL (pH = 8.4), 50 mM KCl |
| Manufacturer of qPCR instrument | StepOnePlus, Thermo Fisher Scientific |
| **qPCR validation** |  |
| Specificity | Analyzed by Melting Curve parameters on each qPCR run and automatic MTP assessment by StepOnePlus software (v2.3). NTC assessment. |
| Method of PCR efficiency calculation | Mean PCR efficiency per amplicon calculated by LingRegPCR program (Ramakers *et al.*, 2003). |
| **Data analysis** |  |
| qPCR analysis program | LinRegPCR program |
| Method of Cq determination | LinRegPCR program |
| Outlier identification | LinRegPCR program |
| Justification of number and choice of reference genes | 2 reference genes tested: UBQ5 (NM_116090) and EF1-α (NM_125432), similarly as performed in (Manacorda *et al.*, 2013) and (Manacorda *et al.*, 2021). For relative quantification of AQPs genes, Bestkeeper algorithm (Pfaffl *et al.*, 2004) was used to obtain a linear combination of both RefGenes. For graphical comparison with Cqs of AQPs, UBQ5 was chosen as the most stable reference gene |
| Description of normalization methods | (Pfaffl *et al.*, 2002)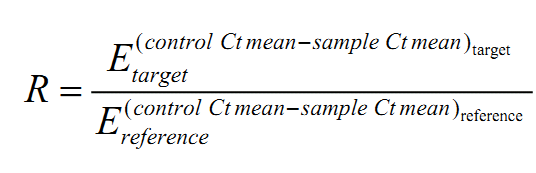 |
| Number of technical replicates | 2 |
| Statistical method | Permutation test |
| Software | fgStatistics software ((Di Rienzo, 2009) (<http://sites.google.com/site/fgStatistics/>) |
| Repeatability (intraassay variation) C_q_ mean SD error | Between 0.05 and 0.55 depending on the assayed amplicon. |

(*) Conditions are described for a generic qPCR assay. Particular annealing/extension temperatures could vary between amplicons. Specific conditions for each amplicon are available upon request.

Further bibliographic resources were the “Getting Started Guide of Applied Biosystems StepOne™ and StepOnePlus™ Real-Time PCR Systems” (Applied Biosystems, 2008).
